# A multicellular developmental program in a close animal relative

**DOI:** 10.1101/2024.03.25.586530

**Authors:** Marine Olivetta, Chandni Bhickta, Nicolas Chiaruttini, John Burns, Omaya Dudin

## Abstract

All animals develop from a single-celled zygote into a complex multicellular organism through a series of precisely orchestrated processes. Despite the remarkable conservation of early embryogenesis across animals, the evolutionary origins of this process remain elusive. By combining time-resolved imaging and transcriptomic profiling, we show that single cells of the ichthyosporean *Chromosphaera perkinsii -* a close relative that diverged from animals approximately 1 billion years ago - undergo symmetry breaking and develop through cleavage divisions to produce a prolonged multicellular colony with distinct co-existing cell types. Our findings about the autonomous developmental program of *C. perkinsii*, hint that such animal-like multicellular development is either much older than previously thought or evolved convergently in ichthyosporeans.

**One-Sentence Summary:** The ichthyosporean *C. perkinsii* develops via symmetry breaking, cleavage divisions, and forms spatially-organized colonies with distinct cell types.

## Introduction

The evolution of multicellular organisms from their unicellular ancestors marks a major transition in the history of life on earth (*1–5*). This transition was accompanied by fundamental developmental challenges such as generating diverse cell types, forming three-dimensional (3D) tissues and establishing overall coordination to drive body plan formation. Asymmetric cell division contributes to cellular diversity (*6–8*), while the formation of 3D tissues relies on the precise coordination of cell divisions, adhesion, and signalling (*9–14*). Compared to other eukaryotic lineages, animal multicellularity stands out by relying on an autonomous developmental program, predominantly driven by intrinsic signals, that drives the emergence of an extensive variety of cell types from a single-celled zygote (*15–17*). Throughout early animal embryogenesis, several ordered processes take place including cleavage divisions, axis establishment, zygotic genome activation and spatial organization of germ layers. This sequence, while displaying remarkable conservation across species, exhibits a degree of developmental plasticity, highlighting the adaptability of development to distinct ecological pressures (*18–20*). Despite this plasticity, the fundamental aspects of this developmental sequence suggest that parts of the underlying program may have originated before the emergence of animals themselves (*21*). This is partly supported by paleontological studies, particularly those focusing on the early Ediacaran Weng’an Biota of the Doushantuo Formation (609 Mya) (*22*), which have uncovered fossils like *Megasphaera*, *Helicoforamina*, *Spiralicellula*, and *Caveasphaera*, which are tentatively classified animal stem groups (*23*). These fossils exhibit morphological features characteristic of animal embryos such as cleavages, Y-shaped cell-cell junctions, tetrahedral 4-cell stages, and a proposed open mitotic strategy (*23–26*). However, their interpretation as stem groups to animals is a subject of ongoing debate(*27*, *28*), as this interpretation strongly relies on the cellular dissimilarities between these fossils and a select few protists, known to be among the closest living relatives of animals.

Animals are closely related to choanoflagellates, filastereans, pluriformeans, and ichthyosporeans (Fig. 1A) (*2*, *29–32*). These lineages not only partly share a genetic toolkit used by animals for development, but can also form transient multicellular structures and display various temporary cell stages in distinct environments (*1*, *30*, *33*). Current evidence indicates that the formation of clonal multicellular choanoflagellate colonies (*34–36*) and the emergence of filastereans aggregates (*37–39*), both morphologically distinct from any known animal embryonic stage or Ediacaran fossil, occur as a facultative response to external chemical cues (*34*, *35*, *40*– *43*). Moreover, while existing results in the choanoflagellate *Salpingoeca rosetta* suggest the presence of morphologically distinct cells within its transient multicellular colonies (*44*, *45*), definitive evidence for a coordinated developmental program driving cell-type differentiation in any close animal relatives remains lacking. These results argue that the animal intrinsic embryonic program orchestrating 3D-cell architecture and spatial cell-type differentiation likely evolved simultaneously with the emergence of animal multicellularity. However, this assumption is largely based on studies focusing on choanoflagellates and filastereans, often overlooking the life cycles, cellular physiology and development of other close relatives.

**Fig. 1:**
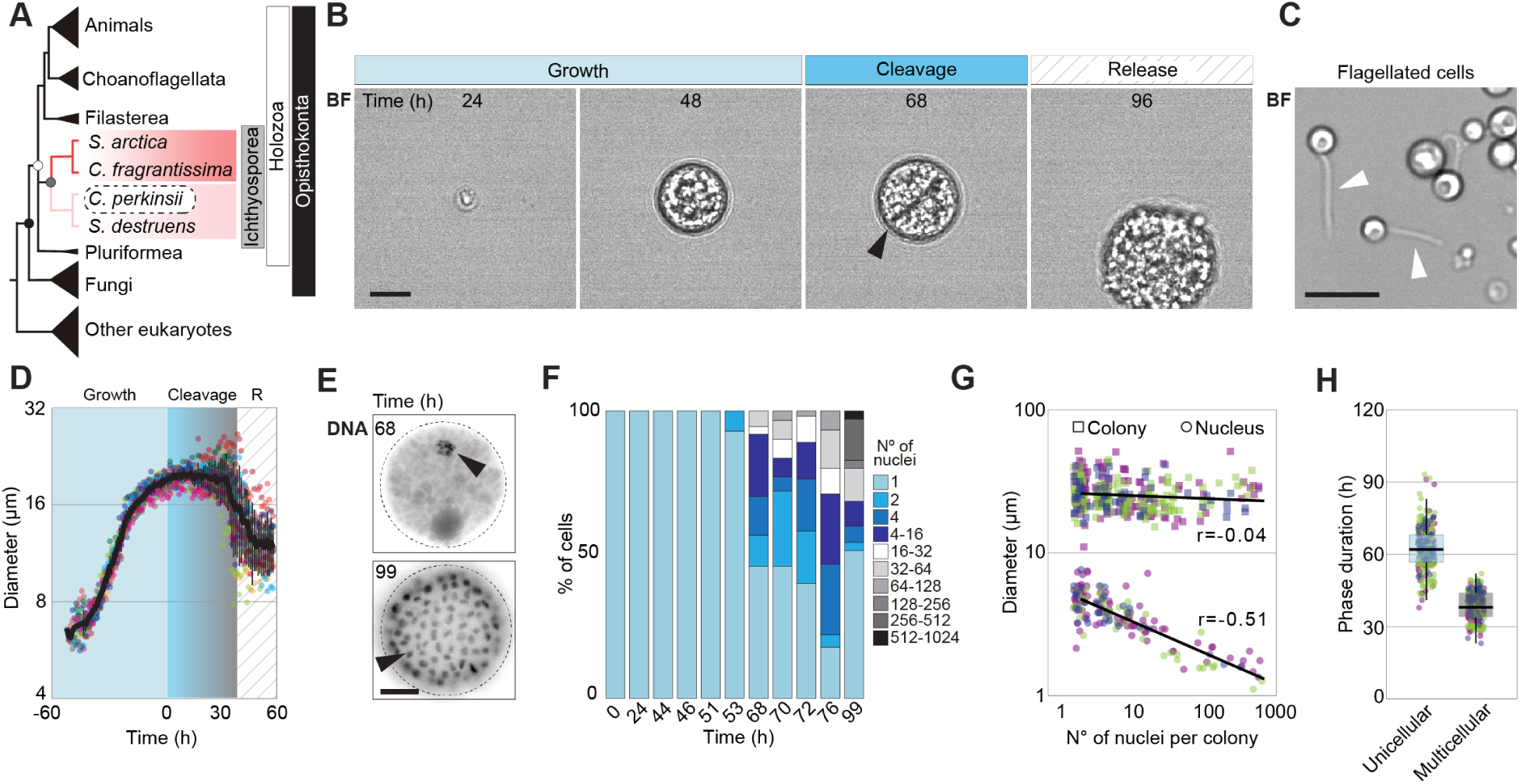
*C. perkinsii* undergoes a series of cleavages at constant volume. **(A)** Cladogram representing the position of ichthyosporeans, featuring ichthyophonids such as *S. arctica* (red) and dermocystids, including *C. perkinsii* (pink), within the eukaryotic tree. **(B)** Time-lapse images of the life-cycle of *C. perkinsii* show cell-size increase, cleavage at the 1-to-2 cell stage (arrow) and release of new-born cells (movie S1). Bar, 10 µm. **(C)** Microscopy of flagellated cells observed following cell release (arrows). Bar, 10 µm. **(D)** Mean cell diameter over time of 10 single cell traces aligned to the 1^st^ cell cleavage reveals the duration of distinct cell stages: Growth, cleavage, and cell release (R). **(E)** Microscopy of DNA-stained cells at 68 and 99 hrs of growth, highlighting the number and size of nuclei at each timepoints (arrows). Bar, 10 µm. **(F)** Distribution of nuclear content of *C. perkinsii* colonies across the life cycle at 23°C measured by microscopy of Hoechst-stained cells (n > 96/time point). **(G)** Distribution of colony (squares) and nucleus (circles) diameters relative to the number of nuclei per colony demonstrates that cellular size remains constant, whereas nuclear size decreases (n = 1571 colony). The colours represent 3 independent replicates. **(H)** Average phase duration for the unicellular and multicellular growth phases (n = 244 cell/colony). The colours represent 3 independent replicates.

Among close animal relatives, the Ichthyosporea, comprising two primary lineages, Dermocystida and Ichthyophonida (Fig. 1A) (*46*, *47*), exhibit a variety of life cycles that combine fungal-like characteristics with transient multicellular stages reminiscent of early animal development (*47*). Most ichthyophonids, including the model *Sphaeroforma arctica*, undergo coenocytic development, characterized by synchronized nuclear divisions without cytokinesis (*48–52*). This process depends on an acentriolar microtubule organizing centre (MTOC) that drives a fungal-like closed mitosis (*53*). Upon reaching a specific nuclear-to-cytoplasmic ratio (*54*), *S. arctica* undergoes an actomyosin-dependent cellularization, during which a transient multicellular layer resembling an animal epithelium is formed before the release of new born cells to repeat the cycle (*50*). In contrast, our recent study reveals that *Chromosphaera perkinsii*, the only cultured free-living dermocystid (*55*), undergoes centriole-mediated, animal-like open mitosis coupled with cell cleavages (*53*). This discovery suggests that *C. perkinsii* proliferates through a palintomic developmental mode, highlighting the vast range of cellular and developmental diversity within the Ichthyosporea. However, nothing is currently known about the spatiotemporal dynamics of the ichthyosporean palintomic life cycle, nor is it clear if it is orchestrated by an autonomous developmental program that exhibits any similarities with early animal embryogenesis.

## Results

To better characterize the development of *C. perkinsii*, we conducted long-term brightfield live imaging of synchronized cell populations (Fig.1B to D, S1A to C and Movie S1). In spite of the light sensitivity of *C. perkinsii* (see methods), our results reveal that synchronized single cells undergo ∼65 hrs of growth until the first cell cleavage (Fig.1, B and D, S1B). Following this initial cleavage, the now multicellular colonies maintain a constant size for an additional 30-hour period, which is followed by the release of hundreds of new born cells (Fig. 1B to D, S1E and Movie S1). Interestingly, released cells were of three types. First, cells without flagella and capable of division, termed “mitotic” (Movie S2). Second, flagellated cells that exhibited flagellar motility termed “flagellates” (Fig. 1C and Movie S3). Third, flagellated cells that remained immobile with dynamic membrane protrusions, and were thus designated “amoeboflagellate” (Movie S2). Counting flagellated cells reveals an increase in their number around the time of cell release (>76 hrs), yet they represent less than 15% of all observed single cells (fig. S1D), indicating that only a fraction of cells differentiates into flagellated cells. These observations suggest that *C. perkinsii* differentiates into at least two distinct cell types, with flagellated cells potentially transitioning between flagellate and amoeboflagellate states.

To gain further insights about *C. perkinsii* development, we fixed and stained the nuclei of synchronized *C. perkinsii* cells throughout their entire lifecycle (Fig. 1E to G). We observe that cells exhibit similar size growth dynamics, though with increased variability, which can be attributed to heterogeneity within the cell population (fig. S1F). Upon counting the number of nuclei per cell, we find that after the first cleavage (∼65 hrs), the nuclear number rapidly increases, surpassing 500 nuclei per colony before cell release (Fig. 1E to G and S1G). As this significant and rapid increase in nuclear content occurs at constant colony size, we note it is accompanied by a gradual reduction in nuclear diameter (Fig. 1E and G). These results show that *C. perkinsii* develops through successive cleavage division while maintaining a constant overall colony volume (Fig. 1G). This multicellular colony stage is maintained for a substantial duration, accounting for approximately 33% of the life cycle in culture (Fig. 1H). In contrast, the epithelium-like multicellular stage in the ichthyosporean model *S. arctica* accounts for only 2% of its life cycle (*50*, *54*).

To better understand whether *C. perkinsii*’s autonomous program is associated with a distinct transcriptional signature, we employed time-resolved transcriptomic profiling. We isolated and sequenced mRNA from two independent synchronized cultures, covering key time points from the single-cell stage through the multicellular stage until cell release (fig. S2A and Table S1). We identify 7,773 transcripts that are differentially expressed (FDR<0.05) across these stages, representing 62% of *C. perkinsii’s* total genes. These genes clustered into five distinct expression patterns throughout development (Fig. 2A, S2A). These patterns highlight the differences between the unicellular phases, both before cleavages (54 hrs) and after release (104 and 120 hrs), compared to the multicellular phase (72 to 96 hrs) (Fig. 2A, S2B). Gene ontology (GO) enrichment analysis reveals that unicellular phase clusters (1, 2 and 5) are enriched in GO terms related to nutrient and lipid biosynthesis and polysaccharide catabolism (Fig. 2B and Tables S2 and 3), whereas multicellular phase clusters (3 and 4) show enrichment in cell division and flagellar differentiation (Fig. 2B). Therefore, prior to the first cleavage, *C. perkinsii* produces and accumulates key cellular components, such as proteins and lipids in a manner reminiscent of oocyte maturation (*56*, *57*). Following the first division, these biosynthetic and catabolic processes are turned off, indicating a gradual consumption of previously synthesized nutrients as the cleavages progress (Fig. 2A and S2A) and only reappear around cell release.

**Fig. 2:**
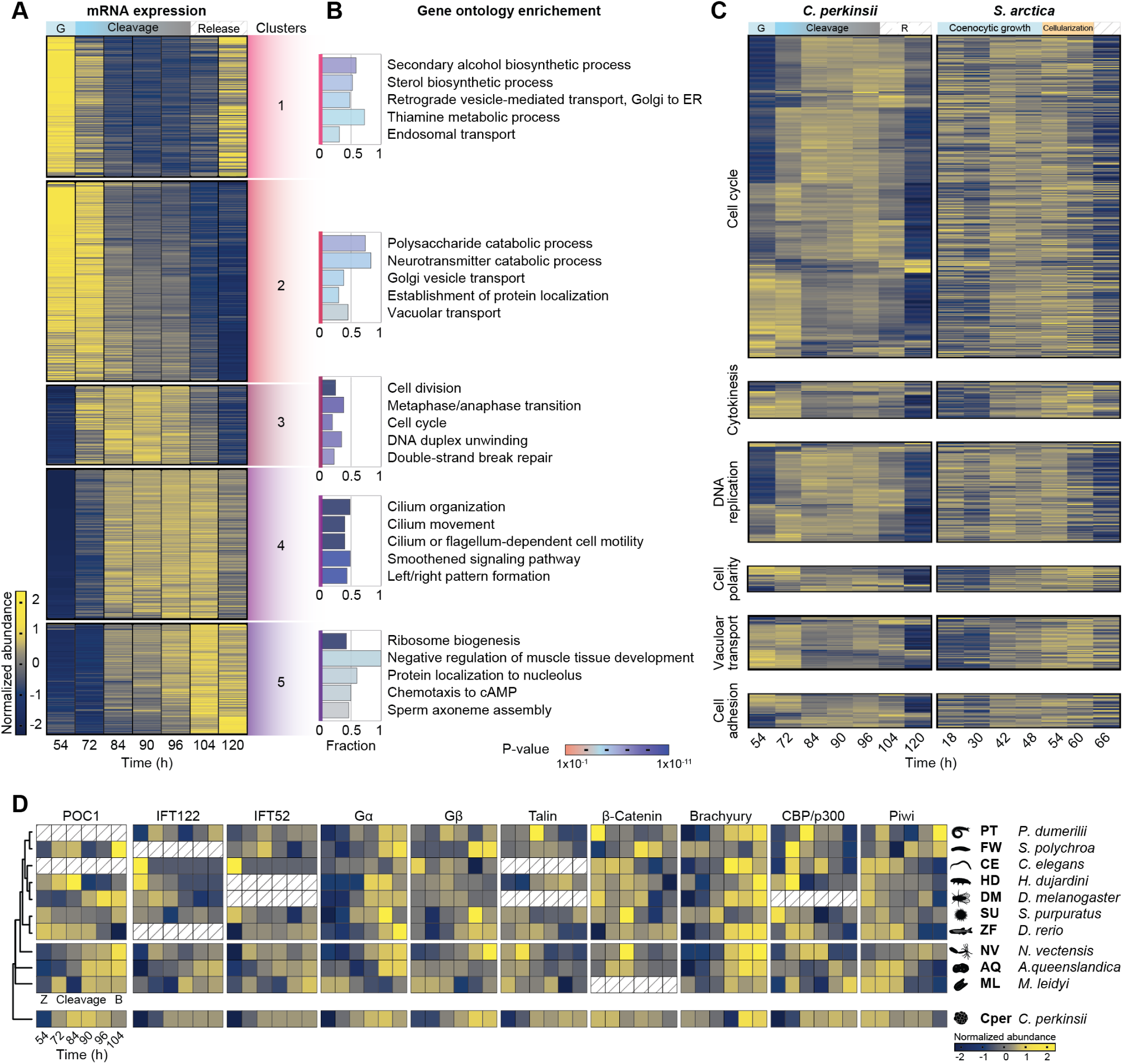
Transcriptional dynamics across the palintomic life cycle of *C. perkinsii*. **(A)** A heatmap of 7773 coding genes clustered into 5 dynamic clusters. **(B)** Gene ontology enrichment of the 5 different dynamic clusters highlighting the patterned transcriptional program of *C. perkinsii* development linked to Tables S2 and 3. **(C)** A comparative heatmap of gene expression dynamics of orthologs between the palintomic *C. perkinsii* and the coenocytic *S. arctica* (*50*) life cycles for key GO categories, including cell cycle (GO:0007049), cytokinesis (GO:0000910), DNA replication (GO:0006260), cell polarity (GO:0007163), Vacuolar transport (GO:0007034), and cell adhesion (GO:0098609). It highlights the distinct transcriptional programs between both ichthyosporean species. The heatmaps for *C. perkinsii* and *S. arctica* have the same row order allowing direct comparison of gene expression patterns of orthologs between the two species. **(D)** A comparative heatmap of gene expression for key regulators involved in flagellar motility (POC1, IFT122, IFT52), cell signalling (Gα, Gβ), adhesion (Talin, β-Catenin), and transcriptional regulation of animal development (Brachyury, CBP/p300, Piwi) dynamics between the *C. perkinsii* (Cper) life cycle and early embryonic development stages of early-branching animals. This comparison spans from the onset of zygote formation (Z) through various cleavage stages and up to the blastula stage (B).

In contrast, the expression of genes driving catabolic processes remains low after release (cluster 2), likely due to nutrient depletion from the medium (Fig. 2A and B, S2B). Towards the end of the multicellular phase a distinct transcriptional pattern emerges (cluster 4) with enrichment in genes for cilia/flagella formation, motility, and left/right patterning in animals (Tables S2 and 3).

To determine if the developmental program of *C. perkinsii* is distinct among ichthyosporeans, we compared it with the transcriptional signature of the ichthyosporean *S. arctica* which undergoes coenocytic growth followed by cellularization. We find that key GO functional categories including the cell cycle, cytokinesis, DNA replication, polarity, and differentiation do not overall correlate between *C. perkinsii* and *S. arctica*, indicating distinct transcriptional programs (Fig. 2C, S2C and D). Despite these differences, certain cytoskeletal regulators previously highlighted and required in *S. arctica* epithelium-like cell layer (*50*) such as formin 2, critical for actin nucleation, correlates between both species regardless of developmental type, suggesting a housekeeping role (fig. S2E). In contrast, formin 5 exhibits an inverse correlation, implying a species-specific role in actin dynamics (fig. S2E). Next, we sought to examine whether the transcriptional program of *C. perkinsii* exhibits similarities with that of early embryonic development in early-branching animals. To accomplish this, we categorized genes from animals and ichthyosporeans into ortho-groups; clusters of genes that originated from a single gene in their last common ancestor. We then analysed the expression patterns of these genes from the onset of zygote formation (Z), through the various cleavage stages, and up to the blastula stage (B) as previously described (*58*) (see methods). By calculating the average correlation of gene expression among orthogroups across time points for *C. perkinsii* and the three earliest-branching animals - the ctenophore *Mnemiopsis leidyi*, the sponge *Amphimedon queenslandica*, and the cnidarian *Nematostella vectensis* - we find an overrepresentation of positively correlated expression patterns indicating similarities in the transcriptional program between *C. perkinsii* and these three early-branching animals (fig. S2G). By focusing on a subset of genes important for flagellar formation (POC1, IFT122, IFT52) (*59*, *60*), cellular signalling (Gα, Gβ, PKAc-α) (*61–63*), cell adhesion (Talin, β-Catenin, Vinculin, Grancalcin) (*64–67*), and transcriptional regulation during animal development (Brachyury, CBP/p300, Piwi, Rbl1, P53, RunX) (*55*, *68–70*), we can observe comparable patterns of gene expression between *C. perkinsii* and basal animals. In certain cases, this expression exhibits conserved temporal patterns across most lineages (Fig. 2D, S2H). Notably, Brachyury, a transcription factor regulating gastrulation, and Piwi, a marker of germline formation, show such conserved expression patterns in many, though not all, animals (Fig. 2D). While we cannot exclude the potential impact of noise from bulk culture asynchrony, ortho-group identification, or developmental time averaging, our results show similar patterns of expression of genes critical for animal embryogenesis during the multicellular development of *C. perkinsii*.

To further explore the development of *C. perkinsii* at the cellular level, we tracked the plasma membrane in synchronized cells using long-term live imaging. Our results show that *C. perkinsii* undergoes a series of ordered cleavages with cortical rotations, starting from a single cell, progressing to a 2-cell stage and then beyond (fig. S3A, Movie S4). Ultimately, this yields a spatially organized multicellular colony surrounded by an external cell wall and occasionally exhibiting a central cavity reminiscent of a blastocoel (fig. S3B).

Due to *C. perkinsii* photo-sensitivity and a need for increased resolution, we opted for imaging synchronized cultures at specific developmental times instead of long-term live imaging. We used three approaches: live-imaging of membrane-stained cells to comprehend overall spatial organisation (Fig. 3A, Movie S5), immunostaining of actin and nuclei to characterize nuclear positioning (Fig. 3B), and ultrastructural expansion microscopy (U-ExM) of microtubules, crucial for antibody accessibility, to reveal spindle orientation during early cleavages (Fig. 3C).

**Fig. 3:**
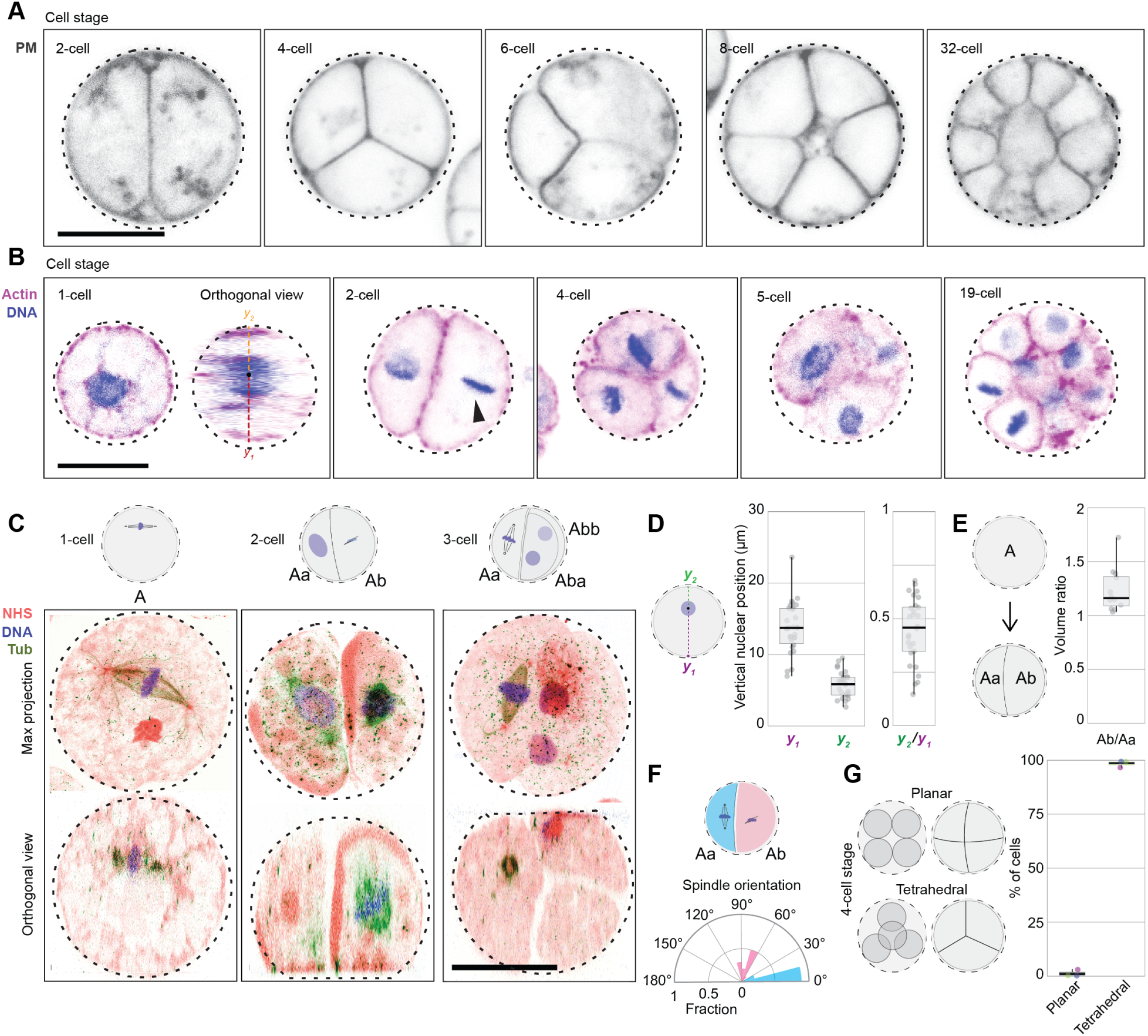
Early developmental patterns in *C. perkinsii*. **(A)** PM-stained live colonies at distinct cell stages, highlighting the patterned cleavages, tetrahedral 4-cell stage and formation of spatially organized multicellular colonies (movie S5). Bar, 10 µm. **(B)** Actin (magenta) and DNA-stained (blue) colonies at distinct cell-stages showcasing the cortical positioning of the nucleus, asymmetrical cell division (in volume and in time), and the formation of a multicellular colony. Bar, 10 µm. **(C)** U-ExM stained colonies for pan-labelling with NHS-Ester (red), microtubules (green), and DNA (blue), highlighting the first mitotic division at the cortex, perpendicular spindles at the 2-cell stage, and a 3-cell stage (movie S6). Bar, 10 µm. **(D)** Box-plot showing the mean vertical localisation of the nucleus at the 1-cell stage prior or at the 1^st^ cleavage highlighting the cortical localisation of the nucleus (n = 28 cells). **(E)** Box-plot illustrating the volumetric ratio following the 1^st^ division resulting in Ab and Aa cells, highlighting the asymmetrical cell division (n =14 colonies). **(F)** A polar plot representing the spindle angle for the Aa and Ab cells during mitosis, demonstrating that the Ab cells divides perpendicular to the 1^st^ cleavage whereas the Aa cells divides parallel to it (n = 11 colonies). **(G)** Box-plot showing the percent of 4-cell stage exhibiting a planar or tetrahedral spatial organisation (n = 108 colonies).

Initially, the nucleus of the 1-cell stage (Cell A) is centrally located, before it migrates to the cortex, where it undergoes mitosis (Fig. 3B to 3D and S3C to S3E). This yields two daughter cells (Aa and Ab), with Ab being ∼1.3 times larger in volume (Fig. 3A, 3E and S3D). Remarkably, we also note that the Ab cell undergoes mitosis ahead of the Aa cell (Fig. 3B, 3C and S3F), resulting in a transient 3-cell stage (Fig. 3C and S3G, Movie S6), thereby highlighting an asynchrony in mitotic entry. Furthermore, we observe that the Ab cell undergoes mitosis perpendicular to the 1^st^ division plane, while the Aa cell undergoes mitosis parallel to it (Fig. 3C and 3F). Therefore, Aa and Ab orient their spindles and divide perpendicularly to each other, ultimately forming a tetrahedral 4-cell stage (Fig. 3B, 3C and 3G). Tracking subsequent cleavages proved more challenging, however we find that the number of cells within the multicellular colonies appears to be variable (Fig. 1F, 1G, and S1G). This potentially suggests that in our experimental setup, *C. perkinsii* undergoes a rotational cleavage pattern before the 8-cell stage, followed by a more chaotic pattern thereafter. Together, these results reveal that the development of multicellular *C. perkinsii* colonies is marked by early symmetry breaking, a phenomenon known to drive cell differentiation across different multicellular systems, followed by formation of a spatially-organized, blastula-like, multicellular colony.

Next, we sought to better understand the differentiation program that results in the formation of distinct cell types, focusing in particular on the flagellated cells, which are easily identifiable using brightfield microscopy (Fig. 1C), expansion microscopy with pan-labelling and antibodies against both beta-tubulin and acetylated tubulin (Fig. 4A and B). As flagellar gene expression starts much earlier than cell release (Fig. 2A to C), we wondered whether the differentiation into flagellated cells occurs post-release, representing a case of temporal cell-type, or whether it differentiates already within the multicellular colony. To this end, we employed expansion microscopy to label flagella and nuclei in the late multicellular colonies (T78, T84, T96). Remarkably, we detect flagellated cells within the multicellular colonies, confirming a differentiation process before cell release (Fig. 4C, Movies S7 and S8). As development progresses and the total number of cells per colony increases, the number of flagellated cells also rises (Fig. 4C and D). However, this number never exceeds 50% of all cells present within the colony, exhibits significant heterogeneity between colonies; moreover, we noted that flagellated cells often cluster spatially within the colony (Fig. 4C to E, Movies S7 and S8). These results suggest the coexistence of at least two distinct cell types within multicellular *C. perkinsii* colonies.

**Fig. 4:**
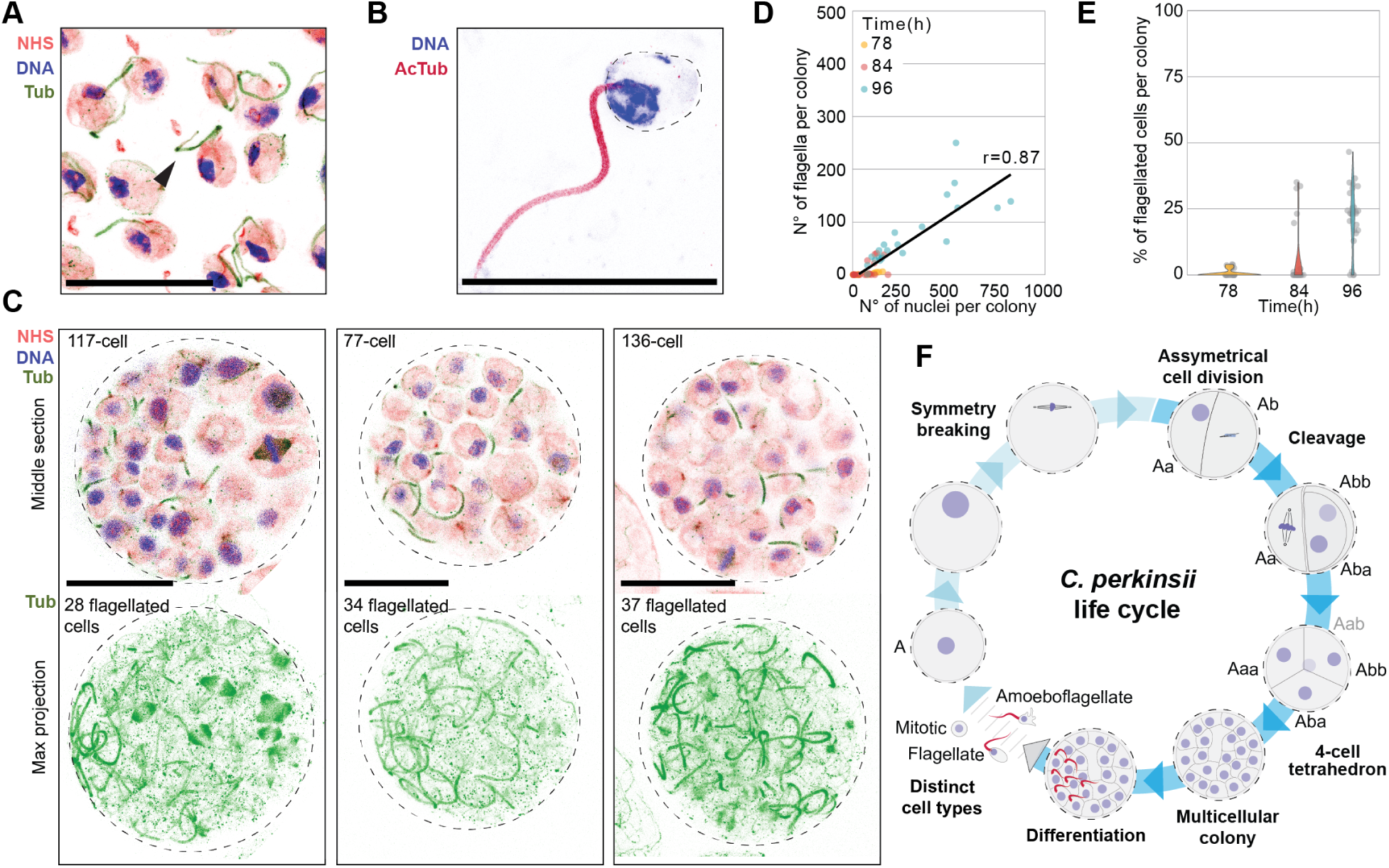
Cell differentiation within *C. perkinsii*’s colonies. **(A)** U-ExM stained cells for pan-labelling with NHS-Ester (red), microtubules (green), and DNA (blue), identifying flagellated cells (arrow). Bar, 10 µm. **(B)** U-ExM stained flagellated cell for Acetylated tubulin (dark red) and DNA (blue). Bar, 10 µm. **(C)** U-ExM stained late colonies for pan-labelling with NHS-Ester (red), microtubules (green), and DNA (blue), highlighting the co-existence of flagellated and non-flagellated cells within the multicellular colony (movies S7 and S8). Bar, 10 µm. **(D)** Dot plot illustrating the number of flagellated cells per colony, which increases with the overall number of cells per colony. Notably, although there is a positive correlation, the increase is not linear, showing that distinct cell types coexist within the multicellular colonies (n > 27 colonies/time point). **(E)** Violin plot showing the percent of flagellated cells within a multicellular colony at distinct developmental times (n > 27 colonies/time point). **(F)** A model representing the developmental program of *C. perkinsii*, beginning with the relocation of the nucleus to the cortex, symmetry breaking at the 1-to-2 cell stage, formation of a tetrahedral 4-cell stage, development into a multicellular colony with intricate 3D architecture, differentiation of a subset of cells into flagellated cells, and release of distinct cell types.

## Discussion

Exploring the evolutionary history that led to the emergence of animal multicellularity has been challenging. This stems notably from the scarcity of fossils predating modern-day animals and the limitations stemming from studying only a few closely related protists, primarily focusing on their genomic content and often neglecting their cellular physiology (*2*, *33*, *71*, *72*). Here, we characterize the life cycle of the ichthyosporean *C. perkinsii*, a close animal relative, providing evidence for an intrinsic and clonal multicellular developmental program. Among all known animal relatives, this program displays unique morphological similarities with early animal embryogenesis. Our results show that *C. perkinsii* undergoes palintomic division (Fig. 4F), involving cell cleavages coupled to open mitosis, diverging from the coenocytic development and cellularization in other studied ichthyosporeans (*52*). Notably, the development of *C. perkinsii* is stereotypical and occurs independently of apparent external trigger, leading to the formation of colonies harboring hundreds of cells (Fig. 4F).

The observation that *C. perkinsii* positions the nucleus cortically in the one-cell stage, undergoes asymmetric cell division, forms spatially organized multicellular colonies, and differentiates into distinct cell types, demonstrates striking morphological and transcriptional parallels with the early stages of animal embryogenesis, which occur following fertilization and precede gastrulation (*20*). This developmental program in *C. perkinsii* challenges the traditional distinctions separating ’simple’ from ’complex’ multicellularity (*5*). ’Complex’ multicellularity is characterized by traits such as cell-cell adhesion, intercellular communication, and a sophisticated developmental program guiding tissue differentiation, traits typically associated with animals, land plants, red and brown algae, Basidiomycota, and Ascomycota. By exhibiting some of these characteristics, *C. perkinsii* emerges as a potential intermediary, blurring the lines between the two classifications.

Although the transcriptional signature of *C. perkinsii*’s development correlates positively with that of early-diverging animals, determining whether it is ancestral to all animals or whether it evolved independently requires still further investigation, including the spatial identification of cell-type signatures at the single cell level. Nonetheless, the ability of *C. perkinsii* to organize into a 4-cell tetrahedral stage and form late blastula-like colonies with clustered flagellated cells within the colony (Fig. 4F), sets *C. perkinsii* apart from previously studied animal relatives including other ichthyosporeans, filastereans and choanoflagellates (*2*, *30*, *33*). Moreover, *C. perkinsii*, which is free-living, exhibits morphological similarities with Ediacaran embryonic-like fossil such as *Megasphaera* (∼600 Mya). These fossils are characterized by Y-shaped cell junctions (*24*), known to require rotating sister cell division planes, flexible cell membranes not limited by separating cell walls, and cell-cell adhesion. While these similarities do not fully elucidate the evolutionary path from Ichthyosporea to animals, they favour the interpretation of *Megasphaera* as an ichthyosporean-like fossil rather than a stem group to animals (*27*).

Altogether, our study highlights the multicellular developmental diversity within the Ichthyosporea, as illustrated by the palintomic program in *C. perkinsii* described here and the coenocytic program of *S. arctica*, reminiscent of Drosophila’s early embryos. With both species representing opposite ends of the developmental spectrum, the Ichthyosporea provide a unique opportunity for a form of "comparative embryology" that can reveal both shared and species- specific mechanisms driving multicellularity at the root of animals.

## Acknowledgments

Pierre Gönczy, Gautam Dey, Alex de Mendoza, Chema Martín-Durán, Hiral Shah, Alexander Woglar for comments on manuscript and general feedback; Hiroshi Suga for *C. perkinsii* cultures; and both the EPFL BioImaging andOptics Platform (BIOP) and the EPFL Gene Expression Core Facility for their support.

## Author contributions

Conceptualization: OD Methodology: MO, CB, NC, JB, OD Investigation: MO, CB, NC, JB, OD Visualization: MO, NC, JB, OD Funding acquisition: JB, OD

Project administration: OD Supervision: OD

Writing – original draft: OD

Writing – review & editing: MO, NC, JB, OD

## Data and materials availability

All data used for quantifications as well as movies are available on figshare (10.6084/m9.figshare.25351078) with the raw images available upon request. The generated code for transcriptomic analysis is accessible on github https://github.com/burnsajohn/Chromosphaera-perkinsii_Transcriptomics and raw RNAseq data are on https://www.ncbi.nlm.nih.gov/sra/PRJNA1091032. Note that the data and code archives are embargoed while the manuscript is under review.

## Supplementary Materials

### Materials and Methods Culture conditions

*C. perkinsii* cultures, previously described in (*53*, *55*), were maintained at 23°C in CpM medium (Yeast extract 3g/L, Malt extract 3g/L, Peptone 5g/L, Glucose 10g/L, NaCl 20g/L) protected from light. Cultures were either streaked onto CpM Agar plates (2% Agar) or grown into liquid CpM medium in rectangular canted neck cell culture flask with vented cap (Falcon; ref: 353108). For maintenance, liquid cultures are propagated monthly from single colonies growing on CpM Agar, and always protected from light. Liquid cultures were refreshed every two weeks (1:1000) dilution and restarted from a cryopreserved stock every 6 months. For synchronization, 10ml of a 6-day-old culture is filtered using a 5µm filter to obtain a homogenous small-celled population which is then diluted 1:100 in CpM medium and tracked for an entire life-cycle (120hrs).

### Microscopy

Imaging of live and fixed *C. perkinsii* cells was conducted utilizing a fully motorized Nikon Ti2- E epifluorescence inverted microscope, which was equipped with a hardware autofocus PFS4 system, a Lumencor SOLA SMII illumination system, and a Hamamatsu ORCA-spark Digital CMOS camera. Imaging was performed using a CFI Plan Fluor 20X objective with a numerical aperture of 0.50, a CFI Plan Fluor 40X Air objective, and a CFI Plan Fluor 60X Oil objective with a numerical aperture ranging from 0.5 to 1.25.

To maintain the temperature at a consistent 23°C, a cooling/heating P Lab-Tek S1 insert (Pecon GmbH) connected to a Lauda Loop 100 circulating water bath was employed. To minimize light-induced phototoxicity, the light source beam was manually adjusted to its lowest setting, and a 715nm LongPass Color Filter (Thorlabs; ref: FGL715S) was utilized to selectively allow far-red wavelengths to pass through. Moreover, we meticulously controlled the imaging frequency and the number of Z-stacks during long-term imaging to prevent cell lysis. Our observations revealed that an imaging interval of 15 minutes, combined with 7 Z-slices over 24 hours, represented the upper limit for optimal imaging conditions, beyond which most cells exhibited abnormal behaviour and lysis. For confocal imaging of membrane-stained, phalloidin-stained and expanded samples, an upright Leica SP8 confocal microscope with an HC PL APO 40X/1.25 Glycerol objective was used.

### Cell fixation and staining

Cell fixation was carried out either using 4% formaldehyde and 250mM Sorbitol for 20 minutes or using pre-chilled Methanol (100%) at -20°C for 7 minutes, followed by two PBS washes. To stain the nuclei, cultures were allowed to settle for 15 minutes at room temperature (RT) before fixation, and Hoechst 33342 nuclear stain (ThermoFisher; 62249) was added at a concentration of 20mM. For actin staining, fixed cells underwent a single PBS wash before adding the F-actin stain Alexa Fluor 488-Phalloidin (ThermoFisher; A12379) at a final concentration of 0.165mM. Before imaging, the fixed and stained cells were concentrated and placed between a slide and coverslip. For live-cell imaging, saturated cultures were diluted 500X in CpM medium inside an 8-well ibidi chamber (Ibidi; ref:80826) or a 35mm dish (Mattek; P35G-1.5-14-C). The plastic cover was removed to ensure oxygenation during the whole experiment period. To reduce evaporation, Silicon oil 100 cSt was added on top (Sigma; 378364). For plasma membrane live staining, FM4-64 (ThermoFisher; T13320) at a final concentration of 10mM from 100X DMSO diluted stock solution was directly added to the medium before imaging.

### Ultrastructural Expansion Microscopy (U-ExM)

U-ExM was adapted from (*53*, *73*, *74*). Initially, cells were fixed in a 4% formaldehyde solution with 250mM Sorbitol for 10 minutes. Following fixation, they underwent two washes with 1X PBS and were subsequently resuspended in 1ml PBS. Coverslips were pre-coated with Poly-l-lysine for an hour, and the fixed cells were then added and allowed to adhere for 1 hour. Anchoring in AA/FA (1% Acrylamide (AA)/ 0.7% Formaldehyde (FA)) was performed for 12 hrs at 37°C. Gelation relied on a monomer solution (19% (wt/wt) sodium acrylate (Chem Cruz, AKSci ref: 7446-81-3), 10% (wt/wt) Acrylamide (Sigma-Aldrich; A4058), 0.1% (wt/wt) N, N’- methylenbisacrylamide (Sigma-Aldrich; M1533) in PBS and was performed at 37°C for 1 hr in a moist chamber. For denaturation, gels were transferred to the denaturation buffer for 15 min at RT and then shifted to 95°C for 1,5 hours. Following denaturation, expansion was performed with water exchanges as previously described(*53*, *73*). Post expansion, gel diameter was measured and used to determine the expansion factor. For microtubule staining of U-ExM gels, mouse primary antibodies targeting beta-tubulin (DSHB; E7) (1/300 dilution) or acetylated-tubulin (Sigma; T6793) and Goat anti-Mouse IgG (H+L) Cross-Adsorbed Secondary Antibodies coupled to Alexa Fluor 488 (Thermofisher; A-11001) (1/500 dilution) were used. For protein pan-labelling, gels were incubated with Alexa Fluor 568 NHS-Ester (Thermofisher; A20003) in NaHCO3 (pH=8.28) for 1.5 hrs. For nuclear staining, Hoechst 33352 was incubated in PBS for 10 minutes before re-expansion in water. For gel mounting, gels were cut to appropriate sizes and attached to pre-coated Poly-l-lysine coverslips and sealed using i-Spacers (Sunjin Lab; #IS013).

### Image analysis

Image analysis was done using ImageJ software (version 1.52) and Imaris (version 10.0.1) (Bitplane). For cell and nuclear diameter measurements (Fig. 1D and G, S1E and S1F), brightfield (live-imaging) and immunostained images were transformed into a binary before using the particle analysis function in ImageJ with a circularity parameter set to 0.15 to 1 to measure the perimeter. As cells are spherical, we computed the cell diameter as D = 2(C/2π). For nuclear content distribution (Fig. 1F and S1G), fixed and Hoechst-stained cells and colonies were imaged and the number of nuclei per cell was either counted using ObjectJ Plugin in ImageJ, or by segmenting the nuclei using the Imaris “surface” tool when the number of nuclei was difficult to count manually. For nuclear position measurements (Fig. 3D) nuclear-stained and U-ExM cells were first oriented with the nucleus positioned at the top. Using the line tool in imageJ we measured the distance between the cortex, either from the bottom (y1) or from the top (y2), and the center of the metaphase plate. For volume ratio measurements of the 2-cell stage (Fig. 3E), we used LimeSeg (*75*) in ImageJ/Fiji (*76*) for semi-automated segmentation of nuclei and phalloidin-stained cells. Briefly, after manual visual inspection, a spherical seed was manually added to the ROI manager for each detected nucleus. These spherical seeds were then inflated with LimeSeg, using two main parameters: 0.8 microns between each surface element and applying a pressure of 0.009 a.u. on the phalloidin channel. Typically, segmentation convergence to the 3D cell shape was achieved in a matter of seconds (Fig. S3D). For spindle orientation measurements (Fig. 3F), U-ExM cells stained for microtubules where the entirety of the spindle, including both spindle poles, were visible were used. The angles were then measured between the plane of the first cleavage and the plane defined by the positions of both spindle poles. To quantify the number of flagellated cells released in the environment (Fig. 4C), we used brightfield or U-ExM images and counted the single cells harbouring a flagellum that were not localized within any multicellular colonies. For the analysis of flagellated cells within multicellular colonies (Fig. 4D and 4E), U-ExM cells stained for Microtubules or flagella (Acetylated Tubulin) and DNA (Hoechst) were examined. Initially, we measured nuclear content using the surface tool in Imaris. Subsequently, flagellated cells were counted using ObjectJ in ImageJ or the surface tool in Imaris, depending on the number of flagellated cells. In instances where there were more than 150 flagellated cells per colony, a manual approach combining the skeletonize tool and ObjectJ in ImageJ was employed. All figures were assembled with Illustrator CC 2020 (Adobe). Several figures were generated using ggplot2 in R version 4.0.5.53.

### RNA isolation, library preparation and sequencing

Synchronized cultures of *C. perkinsii* at 23°C protected from light were sampled at critical time points throughout its entire life cycle (120 hrs). Total RNA was extracted by Trizol and purified using RNA Clean & Concentrator™-5 kit (Zymo; ref: R1013) from 300 to 600ml of culture at each time point. Libraries were prepared using 400ng of RNA using the Illumina stranded mRNA ligation prep (ISML). Libraries were sequenced on NovaS6000 in a PE60 at the EPFL gene expression core facility. We obtained between 51.2 and 104.5 M reads per sample. Reads were trimmed for ISML (Nextera) adapters.

### Gene expression analysis in *C. perkinsii*

RNAseq reads were mapped to the *C. perkinsii* transcriptome (*77*) using Salmon (v1.1.0) (*78*) in mapping-based mode and the full genome (*55*) to generate decoys for accurate read mapping. Salmon quant.sf files were read into the R programming environment (v4.3.1) (*79*) using the tximport (v1.30.0)(*80*) tool. Differential expression analyses within *C. perkinsii* were conducted with edgeR (v4.0.3) (*81–83*). RNA was collected and sequenced for three replicate cultures, however upon examination of the data, one replicate was discarded due to anomalous expression of all time points from that culture as observed in a multiple dimensional scaling plot (fig. S2A), and examination of expression patterns from that culture indicative of a possible infection or contamination of the culture (Table S1). The remaining two replicates (A and C, fig. S2A) were examined for differential gene expression across time by considering differential expression of each time point relative to expression in the first time point using genewise negative binomial generalized linear models with quasi-likelihood tests (glmQLFTest) (*83*) in edgeR. Differentially expressed genes across all time points (false discovery rate (FDR) adjusted p-value < 0.05) were collected for further analyses of expression patterns. Expression patterns were extracted from the differentially expressed genes by fuzzy c-means clustering in R from package e1071 (v1.7.14) (*84*). Considering within-cluster sum of squares for different values of k-clusters, 5 clusters were chosen as the optimal cluster number. Gene expression patterns across time were plotted using the viridis (v0.6.4) (*85*) cividis palette and superheat (v1.0.0) plotting functions (*86*).

### GO functional enrichment analyses

Functional annotation of the *C. perkinsii* transcriptome was conducted by BLASTP (v2.10.0+) (*87*) analysis of *C. perkinsii* predicted peptides against the UniProtKB/Swiss-Prot expertly curated protein database (downloaded Jun 16, 2021, containing 565,254 entries) (*88*). BLASTP hits were filtered by best “homology-derived structure of proteins” (HSSP) distance score (*89*) as in Burns et al. 2016 (*90*). The GO annotations for best Swiss-Prot hits to each *C. perkinsii* peptide were converted to a .goa format file and used for functional enrichment analyses of expression clusters with topGO (v2.54.0) (*91*).

### Comparison of *C. perkinsii* gene expression to *S. arctica* and 10 animal developmental time courses

To compare gene expression patterns across evolutionary divides, we used OrthoFinder (v 2.5.4) (*92*) to cluster genes into orthogroups across ten animals found in Levin et al. 2016 (*58*): *Caenorhabditis elegans*; *Platynereis dumerilii*; *Drosophila melanogaster*; *Strongylocentrotus purpuratus*; *Danio rerio*; *Mnemiopsis leidyi*; *Nematostella vectensis*; *Schmidtea polychroa*; *Hypsibius dujardini*; *Amphimedon queenslandica*; as well as *C. perkinsii*, and *S. arctica* (*50*), and four additional ichtyosporeans: *Amoebidium parasiticum*, *Creolimax fragrantissima*, *Ichthyophonus hoferi*, *Sphaerothecum destruens*; one Corallochytrean: *Corallochytrium limacisporum*; two fungi: *Saccharomyces cerevisiae*, and *Schizosaccharomyces pombe*; and one amoebozoan: *Dictyostelium discoideum*. Transcriptomes and read count data for the ten animals from Levin et al. 2016 (*58*) were downloaded from the associated NCBI GEO database (accession number GSE70185). Transcriptomes were translated using TransDecoder (v5.5.0) (*93*). Peptide files for additional ichthyosporeans and the corallochytrean were downloaded from the Ruiz-Trillo Mulitcellgenome lab resources on FigShare (*77*). The yeast and amoebozoan predicted peptide data (outgroups) were downloaded from their NCBI genome pages (*94–96*). Orthogroups from the predicted peptides of the 20 organisms were predicted using the tool OrthoFinder (*92*).

To compare developmental transcriptional patterns among animals and *C. perkinsii*, we selected timepoints from animal development for each animal that went from the one or two cell stage through the blastula stage, as detailed in Table S4, based on the developmental time courses detailed in Levin et al. 2016 (*58*). We used sliding windows to create an equivalent number of timepoints for each animal’s development, reducing the number of timepoints to match those of *C. perkinsii*. We used the first 6 timepoints of *C. perkinsii* development for comparison with animals (54h, 72h, 84h, 90h, 96h, 104h). The final timepoint (120h), aligning with cell release, was considered to be unique to *C. perkinsii* development and not likely shared with multicellular animals whose multicellular developmental programs continue beyond the blastula stage.

Orthogroups were used for comparing expression patterns across animals. Given that an orthogroup may encompass multiple genes, each exhibiting unique expression patterns, we employed network analysis to segregate these genes based on their expression profiles before conducting cross-species comparisons (fig. S2F). Specifically, we calculated the Euclidean distance between all gene pairs within an orthogroup to quantify expression dissimilarities. This analysis facilitated the construction of a graph where genes, represented as nodes, were interconnected by edges if their expression distance fell below a predefined threshold (threshold of 1.61), thereby indicating similarity. Conversely, genes with distances surpassing this threshold remained unlinked. Subsequently, isolated nodes were eliminated from the graph, which was then subjected to a connected component analysis using the igraph package (version 1.6.0) in R. This step identified clusters of three or more interconnected genes from within or across organisms as distinct expression patterns within the orthogroup, which were then extracted for further analyses. In instances where an organism lacked a gene corresponding to a specific expression pattern within a connected component yet possessed other genes from the same orthogroup, the expression values of these genes were averaged to represent that organism’s contribution to the orthogroup’s overall expression profile. The network procedure allowed analyses of all distinct expression patterns per orthogroup across animals and is illustrated in Fig. 2D and S2H.

Functional annotation of orthogroups was achieved by 1) aligning all proteins within an orthogroup using mafft (v7.487) (*97*); 2) converting that alignment to a hidden markov model using hmmer (v3.3.2) (*98*), and emitting a strict consensus sequence from that hidden markov model. Then 3) using BLASTP of each consensus sequence against the UniProtKB/Swiss-Prot database (downloaded Jun 16, 2021, containing 565,254 entries) (*88*), and collecting the best hit by hssp score as above for *C. perkinsii* genes. The UniProtKB/Swiss-Prot gene names and annotations were transferred to the orthogroups in this way and used for functional inference and GO analyses.

---

**Fig. S1:**
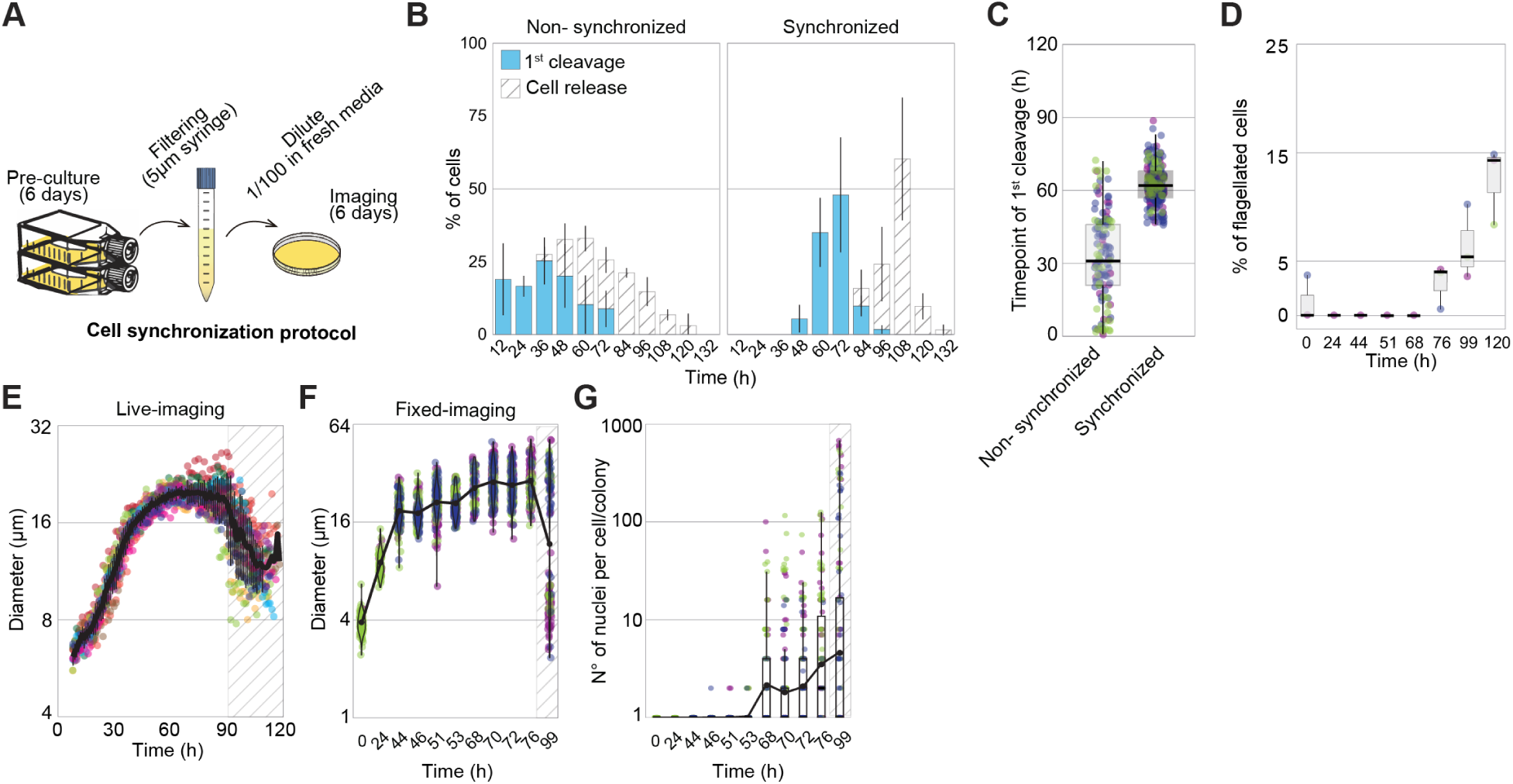
Culture synchronization and growth dynamics of *C. perkinsii*. **(A)** Schematic of the synchronization protocol of *C. perkinsii*. **(B)** Distribution of cells undergoing their first cleavage or cell release throughout the life cycle at 23°C, following either synchronization or not (n > 129 cell/colony per time point). **(C)** The average time at which non-synchronized and synchronized cells undergo their 1^st^ cleavage in a bulk culture (n >129 cell/colony per time point). **(D)** Percentage of free-swimming flagellated cells over time (n > 67 cells/time point). **(E)** Cell diameter over time of 10 single cell traces aligned to time. Variability increases from 75 hrs onwards due to asynchronous cell release. **(F)** Cell diameter over time of fixed cells (n = 1571 cell/colony). The colours represent 3 independent replicates. **(G)** Distribution of number of nuclei per cell/colony across the life-cycle (n = 1571 cell/colony). The colours represent 3 independent replicates.

**Fig. S2:**
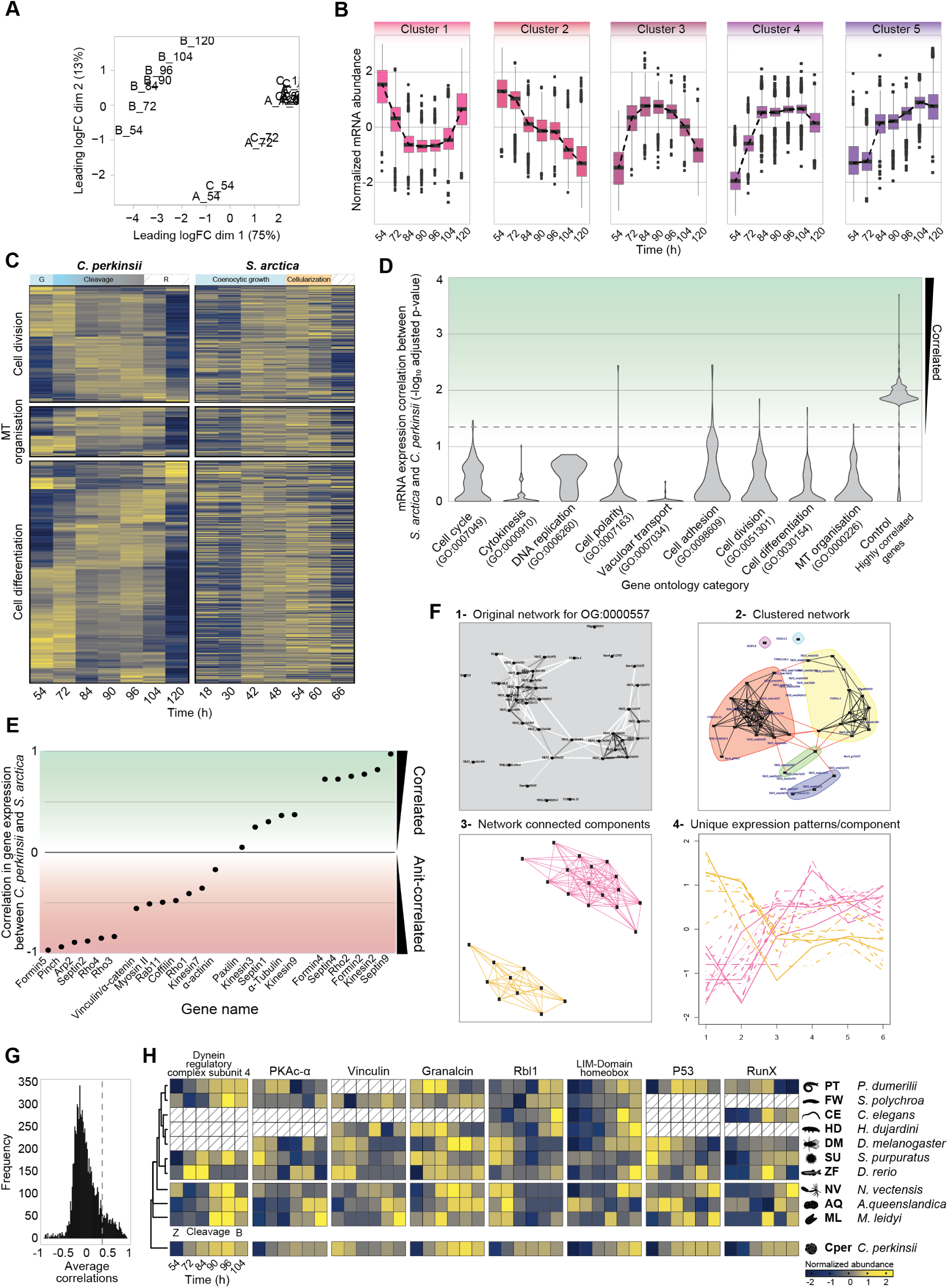
Transcriptional dynamics in *C. perkinsii*’s palintomic lifecycle and comparative analysis with *S. arctica* and early-branching animals. **(A)** Multiple dimensional scaling (MDS) plot of RNAseq counts of three biological replicate developmental time course experiments with *C. perkinsii*. Timepoints for replicates A and C clustered together, while the entire time course of replicate B was set apart indicating markedly different RNAseq counts in all timepoints of replicate B. Differential expression analyses between B, and A and C replicates indicated evidence of an infection in the “B” culture (see Table S1). “B” replicates were discarded for further analyses due to the evidence of infection and outlier expression values. **(B)** Mean expression profile for the 5 distinct gene expression clusters for both replicates. **(C)** A comparative heatmap of gene expression dynamics between *C. perkinsii* and *S. arctica* for key GO categories, including cell division (GO:0051301), microtubule cytoskeleton organisation (GO:0000226), and cell differentiation (GO:0030154). It highlights the distinct transcriptional programs between both ichthyosporean species. **(D)** Statistical evaluation of rowwise correlations between *C. perkinsii* and *S. arctica* for each GO category. For each row, a correlation test was run and the p-values were collected and corrected for multiple testing using the Benjamini-Hochberg procedure. The distribution of adjusted p-values is plotted for each GO category. The horizontal line represents a p-value of 0.05. Genes with a p-value above the p=0.05 line have significantly correlated expression patterns between *C. perkinsii* and *S. arctica.* The plot shows that few genes are correlated for the indicated GO categories. The final category “Control Highly Correlated Genes” shows the p-value distribution when the 300 genes with the highest correlation values between *C. perkinsii* and *S. arctica* are subjected to this analysis–it is the expected distribution if any GO category had genes with highly correlated expression patterns between the two organisms. **(E)** Correlation values of *C. perkinsii* compared to *S. arctica* expression patterns for genes that were determined to be important during the life cycle/development of *S. arctica* in Dudin et al, 2019 (*50*). Values range from strongly anti-correlated to strongly correlated with no discernable functional coherence. **(F)** Demonstration of the network strategy to decouple distinct expression patterns that occur within an orthogroup, across animals. 1) For each OG, a network is constructed with all genes from all organisms within that OG represented as nodes, and edges representing the distance between their developmental expression vectors. 2) The network is subjected to edge-betweenness clustering and only those clusters containing three or more nodes, each with at least three connections, are retained. 3) The resultant clusters can be represented as network connected components. 4) Expression patterns for each connected component are similar within a connected component, but different between connected components indicating separation of distinct expression patterns within an orthogroup. The distinct expression patterns are analyzed separately for each OG across animals for detection of functions with shared gene expression. The procedure of separating distinct expression patterns per OG is more sensitive than the alternative of averaging gene expression within an OG for each animal. **(G)** The distribution of average correlation values of all pairwise correlations between three basal animals (*Mnemiopsis leidyi*. *Nematostella vectensis*, *Amphimedon queenslandica*) and *C. perkinsii* for each orthogroup (gene). Orthogroups with an average correlation value greater than 0.5 (vertical dashed line) were subjected to GO enrichment analyses to determine which biological processes are enriched among genes with highly correlated expression in the development of basal animals and *C. perkinsii*. **(H)** A comparative heatmap of gene expression for key regulators involved in flagellar motility (Dynein regulatory complex subunit 4), cell signalling (PKAc-α), adhesion (Vinculin, Grancalcin), and transcriptional regulation of animal development (Rbl1, LIM-Domain homeobox, P53, RunX) dynamics between the *C. perkinsii* (Cper) life cycle and early embryonic development stages of early-branching animals. This comparison spans from the onset of zygote formation (Z) through various cleavage stages and up to the blastula stage (B).

**Fig. S3:**
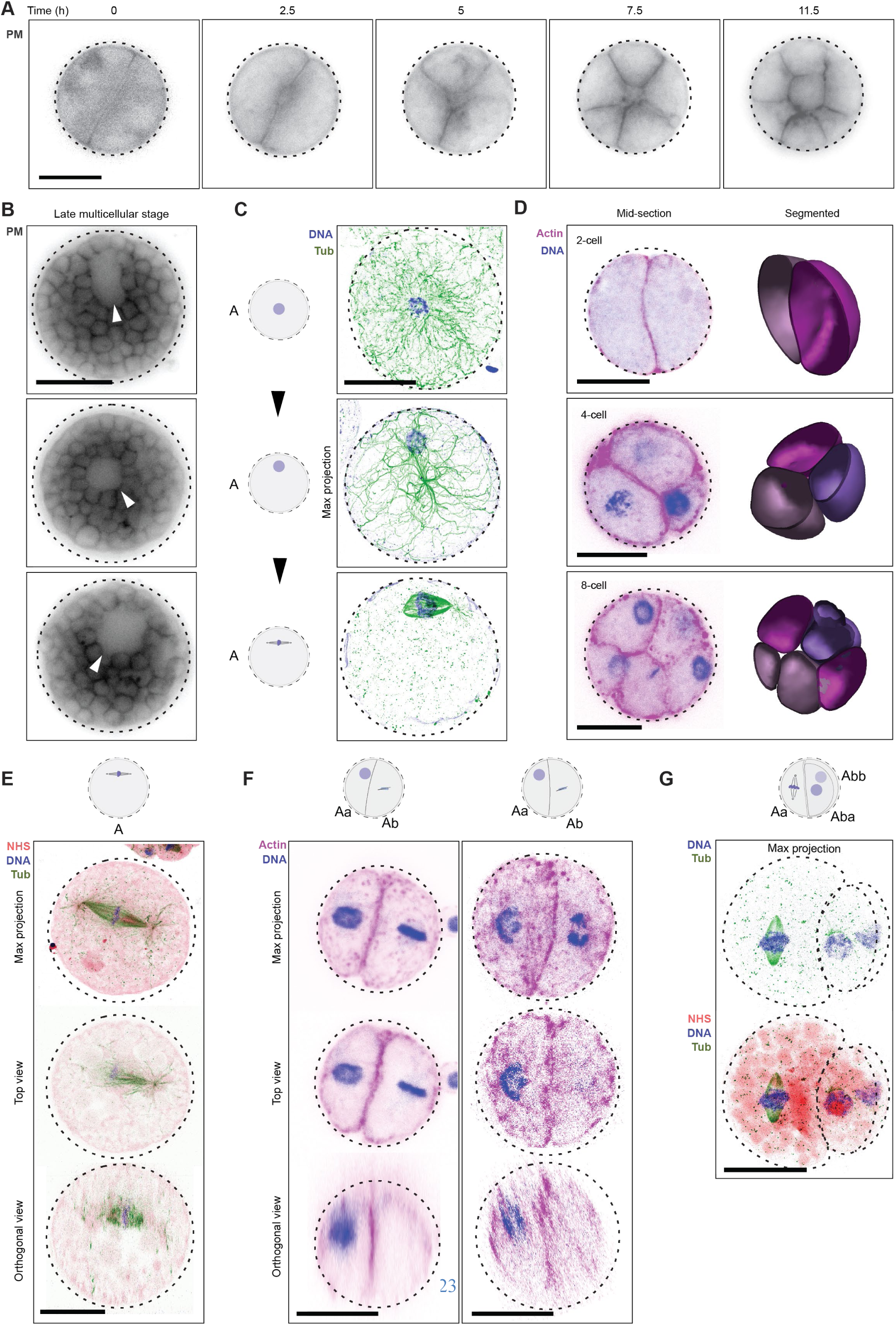
Symmetry breaking during early development of *C. perkinsii*. **(A)** Cleavage dynamics at constant volume over time, visualized using a plasma membrane (PM) marker FM4-64 (movie S4). Bar, 10 µm. **(B)** PM-stained live late colonies exhibiting internal cavities (arrows). Bar, 10 µm. **(C)** U-ExM stained cells for microtubules (green), and DNA (blue), highlighting nuclear migration to the cortex prior to 1^st^ mitotic division. Bar, 10 µm. **(D)** Actin (magenta) and DNA-stained (blue) colonies at various cell stages, accompanied by volumetric segmentation of cells, highlighting the asymmetry in volume following the first cleavage. Scale bar: 10 µm. **(E)** U-ExM stained cell for pan-labelling with NHS-Ester (red), microtubules (green), and DNA (blue), illustrating the first mitotic division at the cortex. Bar, 10 µm. **(F)** Actin (magenta) and DNA-stained (blue) colonies at distinct cell-stages showing the asymmetrical cell division in volume and time between the Aa and Ab cells. Bar, 10 µm. **(G)** U-ExM stained colonies for pan-labelling with NHS-Ester (red), microtubules (green), and DNA (blue), highlighting a three-cell stage. Bar, 10 µm.

## Movies. S1 to S8

**Movie S1:** Time lapse of two synchronized cells of *C. perkinsii* obtained with epifluorescent microscopy. Time interval between frames is 1 hr. The movie is played at 7fps. Both cells can be seen undergoing a full life-cycle with the release of new born cells. Bar, 15 µm.

**Movie S2:** Time lapse of distinct cell types: Ameboflagellate (left) harbouring a flagellum and exhibiting cellular protrusions and a mitotic cell without a flagellum (right) undergoing seemingly cell division. Time interval between frames is 5 sec. The movie is played at 7fps. Bar, 5 µm.

**Movie S3:** Flagellated cells freely moving around following cell release among other non-flagellated cells. Time interval between frames is 1 sec. The movie is played at 7fps. Bar, 10 µm.

**Movie S4:** Time lapse of a PM-stained *C. perkinsii* cell undergoing a series of cleavages at constant volume and showcasing cortical rotations. Time interval between frames is 15 min. The movie is played at 7fps. Bar, 10 µm.

**Movie S5:** Z-stacks of PM-stained live colonies at distinct cell stages, highlighting the patterned cleavages, tetrahedral 4 cell stage and formation of spatially organized multicellular colonies. Bar, 10 µm.

**Movie S6:** Z-stacks of two U-ExM stained colonies for pan-labelling with NHS-Ester (red), microtubules (green), and DNA (blue) at the three-cell stage. Scale bar: 10 µm.

**Movies S7 and S8:** 3D visualisation of U-ExM stained late colonies for pan-labelling with NHS-Ester (grey), microtubules (red), and DNA (cyan), highlighting the co-existence and clustering of flagellated and non-flagellated cells within the multicellular colony. Bar is uncorrected for expansion factor of 4.2X.

---

**Table S1:** Enriched GO terms in differentially expressed genes in *C. perkinsii* replicate B compared to A and C replicates. GO term enrichment is indicative of a viral infection, which explains the distinct RNAseq profile and exclusion of B replicate compared to that of A and C.

**Table S2:** GO term enrichment for the 5 different gene expression clusters throughout the life cycle of *C. perkinsii*.

**Table S3:** *C. perkinsii* genes associated with the GO term enrichment for the 5 different gene expression clusters.

**Table S4:**
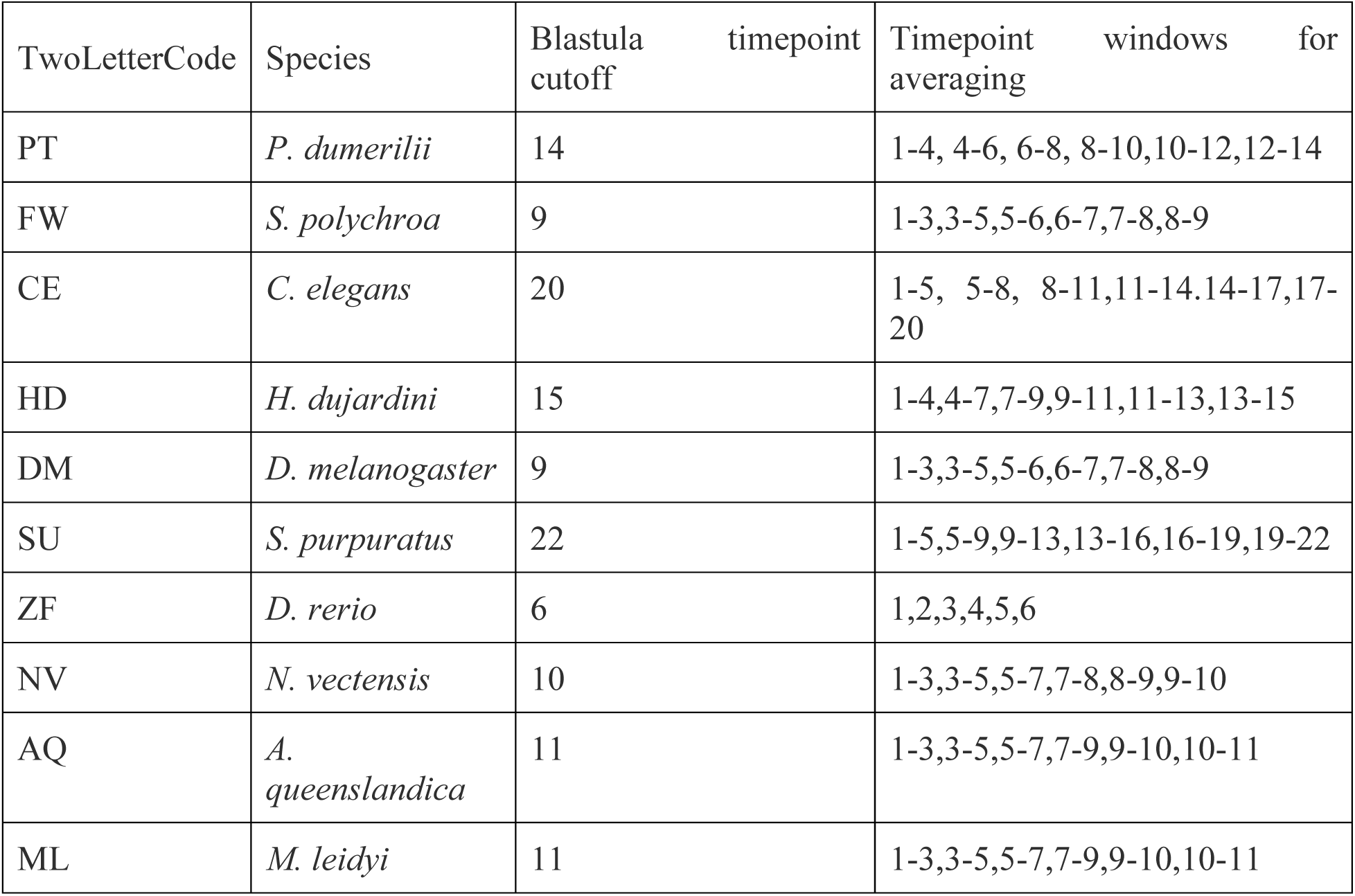
Developmental time points and time averaging windows for animal gene expression comparisons from the one or two cell stage through the blastula stage. The number of time points was selected based on developmental information in Levin et al. 2016(*58*).

